# Family life and cadmium ingestion independently shape offspring microbiomes in a subsocial insect

**DOI:** 10.1101/2024.12.30.630762

**Authors:** Marie-Charlotte Cheutin, Romain Honorio, Joël Meunier

## Abstract

Symbiotic microbes are essential for host health and fitness. In family-living species, these microbes are often acquired through vertical transmission from parents and horizontal transmission from siblings. However, it is unclear how environmental stressors, such as chemical pollution, influence these contributions to the juvenile microbiome. Here, we tested the independent and interactive effects of social environment and cadmium ingestion - a highly toxic and common heavy metal pollutant - on the microbiome of juvenile European earwigs. We reared 900 juveniles either alone, with siblings or with siblings plus the mother. We exposed them to cadmium-enriched food at 0, 25 or 100mg.L^-1^, and analysed their microbiome composition and diversity at the end of the family life period. Our results showed that both social environment and cadmium exposure shaped the juvenile microbial community composition (phylogenetic beta-diversity), with no evidence of an interaction between these factors. In contrast, the microbial local richness (alpha-diversity) remained unaffected by either factor or their interaction. Notably, several specific bacterial taxa, including putatively pathogenic (*Serratia*) and mutualistic (*Lactobacillus*) symbionts, were more abundant in juveniles reared with family members than in those reared in isolation, reflecting classical patterns observed in social species. Overall, our findings suggest that while both social environment and cadmium shape the microbiome of earwig juveniles, family life neither amplifies nor mitigates the effects of chemical exposure. This highlights the robustness of microbial sharing within families, even under strong environmental stress.

## Introduction

Most animals harbour microorganisms that are essential to their health and fitness. These microorganisms play crucial roles in hosts’ biological processes, such as aiding in nutrient acquisition (Berlanga & Guerrero, 2016; Calderón-Cortés et al., 2012), regulating the immune system (Horak et al., 2020), supporting development (Girard et al., 2023; Y. Wang & Rozen, 2017) and providing protection against pathogens (Muhammad et al., 2019; Shukla et al., 2018). The acquisition of these microorganisms, whether from the environment or from parents (*e.g.,* transovarial pathway), is thus essential for the host, especially for juveniles that are born or hatched with an underdeveloped microbiome (Videvall et al., 2019) or that have lost a large part of their microbiome during development, such as through moulting in insects (Cheutin, Boucicot, et al., 2024; Manthey et al., 2022; Schmidt & Engel, 2021).

In group-living species, high nest fidelity and high frequency of social interactions between group members are often key factors facilitating the acquisition of core microorganisms by juveniles (Onchuru et al., 2018). In family groups, this acquisition occur through two main pathways. The first pathway is vertical transmission through parental care - a widespread phenomenon across animal taxa (Trumbo, 2012). Parental care typically allows the transfer of microorganisms and community of microorganisms from parents to offspring through behaviours such as trophallaxis and coprophagy (Gurevich et al., 2020; Hosokawa et al., 2012). The second pathway involves horizontal transmission between juveniles. This may occur through active and cooperative behaviours among siblings, such as stomodeal (mouth-to-mouth) or proctodeal (mouth-to-anus) trophallaxis, that facilitate direct microbial transfer between individuals, or through passive processes involving the excretion and consumption of feces by family members (Lombardo, 2008; Onchuru et al., 2018; Powell et al., 2014). Together these vertical and horizontal pathways are important drivers of the juvenile microbiome (Onchuru et al., 2018). However, the relative contributions of each pathway during family life, and the influence of external factors on these contributions, remain largely unexplored.

Multiple external factors can directly or indirectly shape the vertical and horizontal transfers of microorganisms within families, whether by directly modifying the composition of the microbiome of donor individuals and/or the ability of parents and siblings to perform these transfers (Bright & Bulgheresi, 2010; Hosokawa & Fukatsu, 2020; Sachs et al., 2011). Among these factors, human-induced chemical pollution, particularly heavy metals, has gained increasing attention due to its widespread impact on ecosystems. Heavy metals accumulate in soil, water, and air as a result of agricultural activities, urbanization, and industrial processes (Johnson & Munshi-South, 2017; Seto et al., 2012). They are known to disrupt microbial communities across different environments (Mogren & Trumble, 2010; Sun et al., 2020; S. Zhang et al., 2015). For example, heavy metals (such as cadmium, lead or mercury) disrupt soil microbial communities by reducing bacterial richness and suppressing beneficial taxa (such as *Pseudomonas* and *Bacillus*) which are essential for plant health and nutrient cycling (Z. Cheng et al., 2023; Q. Wang et al., 2022). Similarly, in animal hosts, heavy metal exposure disrupts gut microbial composition, leading to shifts in community structure and metabolic functions. For instance, on earthworms, cadmium exposure affects key taxa (such as *Verrucomicrobia* and *Firmicutes*) which play important roles in digestion and detoxification processes (Wu et al., 2020).

Among heavy metals, cadmium is one of the most biotoxic and mobile elements in ecosystems (Kubier et al., 2019; Luo et al., 2020; Mulligan et al., 2001). Cadmium alters the behaviour, cognition, development, survival or fecundity of many organisms ranging from mammals to insects (*e.g.,* Arito et al., 1981; Gustin et al., 2018; Honorio et al., 2021). Research on cadmium’s effects on animal microbiomes has expanded only recently, particularly regarding its influence on insect-associated microbial communities. Studies on honeybees (*Apis mellifera*) and pygmy grasshoppers (*Tetrix tenuicornis*) have shown that cadmium exposure alters their gut microbiomes, increasing the abundance of opportunistic pathogens such as *Aeromonas* and *Chryseobacterium*, possibly reducing host immune responses and fitness (Li et al., 2021; Rothman et al., 2019). In contrast, some gut microbes appear to confer resistance to cadmium toxicity. For instance, in mice, specific strains of *Lactobacillus plantarum* can sequester cadmium in the intestine, acting as a protective barrier against its toxic effects when administered as probiotics (Zhai et al., 2014, 2016). Despite these insights, the broader implications of cadmium-induced microbiome disruptions remain poorly understood, particularly in terms of microbial transmission between family members. If cadmium alters the gut microbial composition of a host, it may thus disrupt both vertical and horizontal transfer of symbiotic bacteria, potentially leading to shifts in community composition, reduced microbial richness, and consequently major impairments in host digestion, immunity, and development (Teyssier et al., 2018, 2020).

In this study, we investigated how the social environment and cadmium ingestion influence the whole microbiome (encompassing gut, internal non-gut, and surface bacteria) of European earwig juveniles (*Forficula auricularia*). The European earwig is a hemimetabolous insect, meaning that juveniles (called nymphs) and adults share similar morphology and ecology. Nymphs undergo five moults before reaching adulthood (Honorio et al., 2024; Tourneur et al., 2020). Mothers live with their young nymphs in family groups that persist for about two weeks after egg hatching. During this period, mothers frequently interact with their nymphs (Kölliker, 2007; Kölliker & Vancassel, 2007), engaging in grooming, trophallaxis and coprophagy (Falk et al., 2014), which are behaviours typically known to facilitate microbial transfer among social insects – unlike solitary species that primarily acquire their microbiome from a internal route or from their food environment (Onchuru et al., 2018; Y. Wang & Rozen, 2017). However, family life is facultative in the European earwig, as nymphs quickly become mobile after hatching, can forage for themselves, and can reach adulthood without parents and siblings (Falk et al., 2014; Meunier & Kölliker, 2012; Van Meyel & Meunier, 2022). Recent studies indicate that the earwig microbiome is predominantly composed of Proteobacteria and Firmicutes, with dominant bacterial families such as Lachnospiraceae, Enterobacteriaceae or Pseudomonadaceae varying among individuals (Cheutin, Leclerc, et al., 2024; Van Meyel et al., 2021). This microbial composition can be altered by social and chemical stresses, including the absence of mothers during early life stages and the consumption of the antibiotic rifampicin (Cheutin, Boucicot, et al., 2024; Greer et al., 2020; Van Meyel et al., 2021). In contrast, cadmium exposure appears to have minimal effects on the behaviour, survival and development of both adults and juveniles in the European earwig (Honorio, Depierrefixe, et al., 2023; Honorio, Moreau, et al., 2023), which stands in contrast to the typical negative effects observed in most animal species (*e.g.,* Arito et al., 1981; Gustin et al., 2018; Honorio et al., 2021). Here, we investigated whether cadmium exposure can alter the microbiome of the European earwig, and whether the presence of siblings during family life may mitigate these potential alterations, thereby explaining the apparent absence of the typical cadmium-induced phenotypic effects observed in other species.

We reared 900 earwig juveniles either isolated (alone), with siblings or in a family group (with siblings and the mother) and exposed them to one of three concentrations of cadmium (0, 25 and 100 mg.L^-1^) for the entire duration of family life, *i.e.* 14 days (Meunier, 2024). We then extracted the whole microbiome from a subset of 90 nymphs (10 per combination of social environment and cadmium concentration) and used 16S rRNA metabarcoding to analyse their alpha- and beta-diversity, as well as identify potential treatment-specific bacterial taxa. If the frequent grooming, trophallaxis and allo-coprophagy behaviours expressed among siblings and between mother and nymphs mediate microbial (horizontal and vertical, respectively) transfers to juveniles, we predict greater microbial (alpha and/or beta) diversity in nymphs living in group (sibling and family groups) compared to isolated nymphs. Additionally, if mothers possess microbial taxa that are not environmentally accessible to the nymphs, we expect that some microbial taxa will be specific to nymphs reared with their mother. If cadmium alter the microbiome composition of the hosts, we predict a change in microbial diversity in nymphs treated with cadmium. This change could be manifested as a reduced diversity due to the elimination/selection of certain taxa, or as an increased diversity related with a “dysbiosis” by the proliferation of non-core members (*i.e.,* Anna Karenina principle, Zaneveld et al., 2017). Finally, if cadmium ingestion alters the social transmission of the microbiome to the nymphs, we predict that the nymphs’ microbiota will be shaped by an interaction between both parameters: for instance, the diversity of the nymphs’ microbiome would depend on the presence or absence of siblings and mothers only in the absence of cadmium exposure (or vice versa).

## Material and methods

### Laboratory rearing and experimental social environment

This study involved the offspring of 30 females collected in a peach orchard near Valence, France (Lat 44.9772790, Long 4.9286990). All these individuals belonged to *Forficula auricularia* Linneaus, 1758, also called *Forficula auricularia* clade A (González-Miguéns et al., 2020). Just after collection, these females were reared under standard laboratory conditions until oviposition and egg hatching (Meunier et al., 2012). In brief, we maintained the group of females with males for four months to allow uncontrolled mating (Sandrin et al., 2015) and then isolated each mated female in individual containers to allow egg production. This isolation occurred at the end of October, approximately 50 days before the typical oviposition period for this species. From the isolation date, each female received 50 mg of a standard laboratory-made diet (*i.e.,* mostly composed of pollen, cat food, carrots, agar (see details in Kramer et al., (2015) without preservative acids – not useful in this study), covered by one of three concentrations of cadmium (see details below). The food (including cadmium) was changed twice a week until oviposition. At that time, we removed the food as mothers typically stop their foraging activity to care for their eggs (which typically lasts for about 50 days; Kölliker, 2007).

One day after egg hatching, we randomly assigned 30 nymphs per family (mean clutch size per family ± SD = 51.6 ± 12.7 nymphs) to three experimental groups: family group (10 nymphs), sibling group (10 nymphs), and isolated nymphs (10 nymphs). The family group consisted of 10 nymphs reared with their mother, the sibling group consisted of 10 nymphs reared together without their mother, and the isolated group consisted of single nymphs reared in isolation. This setup allowed us to compare the microbiome of isolated nymphs with that of nymphs maintained in groups, either with siblings only or with siblings and their mother. We then provided each experimental group with 50 mg of the standard laboratory-made diet described above, mixed with one of the three concentrations of cadmium. Each mother and her nymphs received the same cadmium treatment throughout the experiment. We maintained these experimental groups in Petri dishes lined with moistened sand at 20-18°C under a 12:12h light:dark cycle. To standardise the space available per individual, we maintained the family and sibling groups in large Petri dishes (diameter 9 cm, height 2 cm), and the isolated nymphs in small Petri dishes (diameter 3.6 cm, height 1 cm). Fourteen days later, which marks the end of family life under natural conditions (Meunier et al., 2012), one nymph per experimental group and cadmium concentration was transferred to an Eppendorf tube and stored at −80°C until the extraction of the genomic material. Hence, a total of 30 nymphs were obtained for each concentration of cadmium, comprising 10 that were raised in a family group, 10 that were raised with siblings only, and 10 that were raised in isolation. This experiment ended at this point.

### Cadmium exposure

We exposed the 30 mothers and their nymphs to cadmium using the protocol and concentrations presented in Honorio, Moreau, et al., (2023). In particular, we applied 10 μL drops of each cadmium solution directly onto the 50 mg of food provided to individuals. This quantity and application method ensured that the food piece was homogeneously covered by the solution. We used cadmium concentrations of either 0, 25, or 100 mg.L^-1^ (corresponding to 0, 5, 20 mg.kg^-1^ of food, respectively), which were obtained by diluting cadmium chloride powder (Sigma-Aldrich #202908, CAS number: 10108-64-2, purity: 99.99%) in MilliQ water. We chose cadmium exposure *via* ingestion because this is the main source of cadmium exposure in nature (H. Zhang & Reynolds, 2019). We chose these concentrations of aqueous solutions to cover both those found in the food chain and soil (Butt et al., 2018; FitzGerald & Roth, 2015; H. Zhang & Reynolds, 2019) and those known to induce behavioural and microbial changes after ingestion in other species (Dési et al., 1998; Rothman et al., 2019; Su et al., 2021). A previous study confirmed that this method of exposure ensures the uptake of cadmium by earwig females (Honorio et al., 2025; Honorio, Moreau, et al., 2023).

### Genomic extraction and 16S amplification

The 90 nymphs were individually isolated in 2mL Eppendorf tube and immediately stored at −80°C until the extraction of both external and internal DNA material (*i.e.,* whole individual body) using the NucleoMag® Tissue extraction kit (Macherey-Nagel™, Düren, Germany). We then amplified the hypervariable region V3-V4 of the 16S rRNA gene using the primers 343F (5′ - ACGGRAGGCAGCAG - 3′) and 784R (5′-TACCAGGGTATCTAATC – 3′) (Muyzer et al., 1993) coupled with platform-specific Illumina linkers. PCR reactions were processed using the Taq Polymerase Phusion® High-Fidelity PCR Master Mix with GC buffer following the manufacturer’s instructions (Qiagen, Hilder, Germany). Each PCR reaction involved an initial step of 30 sec denaturation at 98°C, followed by 22 amplification cycles (10 sec denaturation at 98°C; 30 sec annealing at 61.5°C; 30 sec extension at 72°C), and ended with a final step of 10 min extension at 72°C. Each reaction was checked for the probe specificity and amplification success with an electrophoresis migration. Extraction and amplification steps involved several blank controls to confirm that samples were not contaminated by environmental microorganisms. As blanks were all negative, we did not send them for sequencing (Figure S1). Samples were amplified in duplicate and equally pooled for a final product of 30 µL which was further sequenced using 2×250bp Illumina MiSeq technology at the Bio-Environnement platform (University of Perpignan, France). Finally, two of the 90 nymphs used for genomic extraction were removed from the study as one did not amplify (isolated nymphs) and one did not respect the experimental condition (the sibling group was only constituted by two nymphs instead of 10). Both samples were exposed to 100 mg.L^-1^ Cd.

### Bioinformatic process

We used the DADA2 algorithm v1.24.0 (Callahan et al., 2016) in R software v4.2.0 (R Core Team, 2024) to process sequences. Libraries were first trimmed for primers and reads were truncated at 240 base pairs (both forward and reverse), filtered with their quality profiles, cleaned, dereplicated and inferred into Amplicon Sequence Variants (ASVs) (Glassman & Martiny, 2018). The resulting chimeras were removed and taxonomic assignment using the SILVA reference database (release 138) (Quast et al., 2012). Sequences were aligned using the MAFFT program (Katoh, 2002) and entered into a phyloseq object using the *phyloseq* package v1.40.0 (McMurdie & Holmes, 2013). We inferred the phylogenetic tree associated with these sequences using the FastTree 2 tool (Price et al., 2010) and the *Phangorn* package v2.8.1 (Schliep, 2011). We removed 2 211 sequences that belonged to 65 ASVs that were assigned as mitochondrial sequences or not assigned at least at the Phylum level. We also filtered the sequence using a standard method that (1) preserves rare sequences while filtering out statistically random sequences that may introduce noise from exaggerated inter-individual variability (*i.e.,* unique sequences present in only few individuals), and (2) avoids relying on subjective or arbitrary occurrence and abundance thresholds (Magurran & Henderson, 2003). Specifically, we calculated an index of dispersion (*i.e.,* the variance to mean ratio, VMR) for each ASV. We then tested whether these indices followed a Poisson distribution, checking if they fall within the 2.5 and 97.5% confidence limits of the Chi^2^ distribution (Krebs, 1999) and we preserved the ASVs with index values outside these confidence limits (Figure S2). This removal represents 1.5% of the sequences in the initial dataset, for an occurrence that did not exceed three individuals out of 90.

### Microbial diversity and indicators

We tested the effects of social environment and cadmium ingestion on the alpha- and beta-diversity of the resulting microbiome of each host nymph. To calculate alpha-diversity, we first rarefied all samples to 11 637 sequences corresponding to the threshold at which rarefaction curves reach their stationary phases (Cameron et al., 2021; Figure S3). The rarefaction involved the removal of one sample of family nymphs (100 mg.L-^1^ of Cd), which contained fewer reads than the threshold. We then calculated the observed richness (*i.e.,* ASV or bacterial genus counts), the Shannon entropy that integrates the abundance of each ASV (or genus) present in each samples, and their respective equivalents the Faith and Allen indices that integrate the phylogenetic relationships between ASVs (or genus). For beta-diversity, we first transformed the community matrix in relative abundance then calculated the Jaccard distance between samples with ASVs (or genus) presence/absence and its quantitative equivalent Bray-Curtis. As for alpha-diversity, we also used their phylogenetic equivalents with the Unifrac metrics for qualitative (unweighted) and quantitative (weighted) distances (Chao et al., 2010; Yang et al., 2021). The dissimilarities between samples were projected into a Principal Coordinates Analysis (PCoA), which we plotted on the first two axes. The significant microbial genera correlated along these axes were projected through the PCoA using the function *envfit()* from the package *vegan* v2.6-4 (Oksanen et al., 2022). Alpha- and beta-diversity were also calculated at genus level by merging all ASVs in their belonging genus using the function *tax_glom()* from the *phyloseq* package (McMurdie & Holmes, 2013).

### Statistics

All statistical analyses were performed using R v4.3.4 (R Core Team, 2024). First, we analysed the alpha-diversity of nymph microbiome using four linear mixed models (LMER) with the *lme4* v1.1-31 (Bates et al., 2015) and analyses of variances (Anova) were run using *car* version packages v3.1-1 (Fox & Weisberg, 2019). In these models, we entered either the observed richness, Shannon entropy, Faith or Allen indices as a response variable, and social environment (isolated, sibling or family nymphs) and cadmium concentration (discrete; 0, 25 or 100 mg.L^-1^) as explanatory variables. We also included the interaction between these two factors to test whether cadmium ingestion affects the social transmission, or wether being in a specific social context modulates the effect of the cadmium ingestion on the microbiome. To control for the same origin of each social environment, we also entered the mother identity as a random effect in each model. When necessary, we conducted post-hoc pairwise comparisons using the estimated marginal means via the *emmeans* package v1.8.0 (Lenth, 2022). The R^2^ corresponding to each pairwise comparison was calculated with the *MuMIn* package v1.47.1 (Bartoń, 2022). Second, we analysed the beta-diversity of nymph microbiome using four permutational analyses of variances (PERMANOVAs). In these models, we entered either Jaccard, Bray-Curtis, unweighted and weighted Unifrac metrics as response variables, and social environment, cadmium concentration and their interaction as explanatory variables. We specified the origin of the mother as strata to control for the effects of this variable. We conducted post-hoc pairwise comparisons using the *pairwiseAdonis* package v0.4.1 (Martinez Arbizu, 2017). Overall, we ran these eight models on diversity indices calculated on both ASVs and Genus levels. All models were simplified after AICc comparisons by removing the non-significant interactions. Finally, to identify bacterial indicators of both the social environment and the cadmium concentration, we used the *DESeq2* package v1.36-0 (Love et al., 2014). Each raw counts were clustered into bacterial genera, normalized by its geometric mean, and then compared to a negative binomial distribution. The estimated coefficient (Log2FoldChange) for each genus in each sample group was transformed into a Z-statistic, compared to a normal distribution, and then corrected for multiple comparisons.

We checked all model assumptions using the *DHARMa* package (Hartig, 2022) and the homoscedasticity of the variances between groups of each variable with the function *betadisper* from the *vegan* package v2.6-4 (Oksanen et al., 2022). To fulfil model assumptions, we log-transformed alpha proxies (*i.e.,* Observed richness, Shannon, Faith and Allen indices) as response variable. All p-values were adjusted for multiple testing with False Discovery Rate method (FDR) and Benjamini-Hochberg correction (Haynes, 2013).

## Results

### Overall microbial composition of the nymphs

Overall, the microbiome of the tested nymphs included 2 075 719 sequences (98.52% abundance of the initial dataset) representing 590 different ASVs (31.45% richness of the initial dataset; Figure S2). While Proteobacteria largely dominated the microbiome (78.7%), the Bacteroidota represented 15.2%, and the Firmicutes 5.43%. Finally, the microbiome also contained other bacterial phyla at a low level, such as Actinobacteriota, Bdellovibrionota and Patescibacteria (all < 1 %). The dominant genera (Figure 1) were *Buttiauxella* (Proteobacterales, Enterobacteriacae) (32.1 % of all reads), *Chryseobacterium* (Bacteroidota, Weeksellaceae) (11.5%), *Acinetobacter* (Proteobacteria, Moraxellaceae) (6.33%), *Serratia* (Proteobacteria, Yersiniaceae) (6.31%) and *Pseudomonas* (Proteobacteria, Pseudomonadaceae) (5.34%).

**Figure 1.**
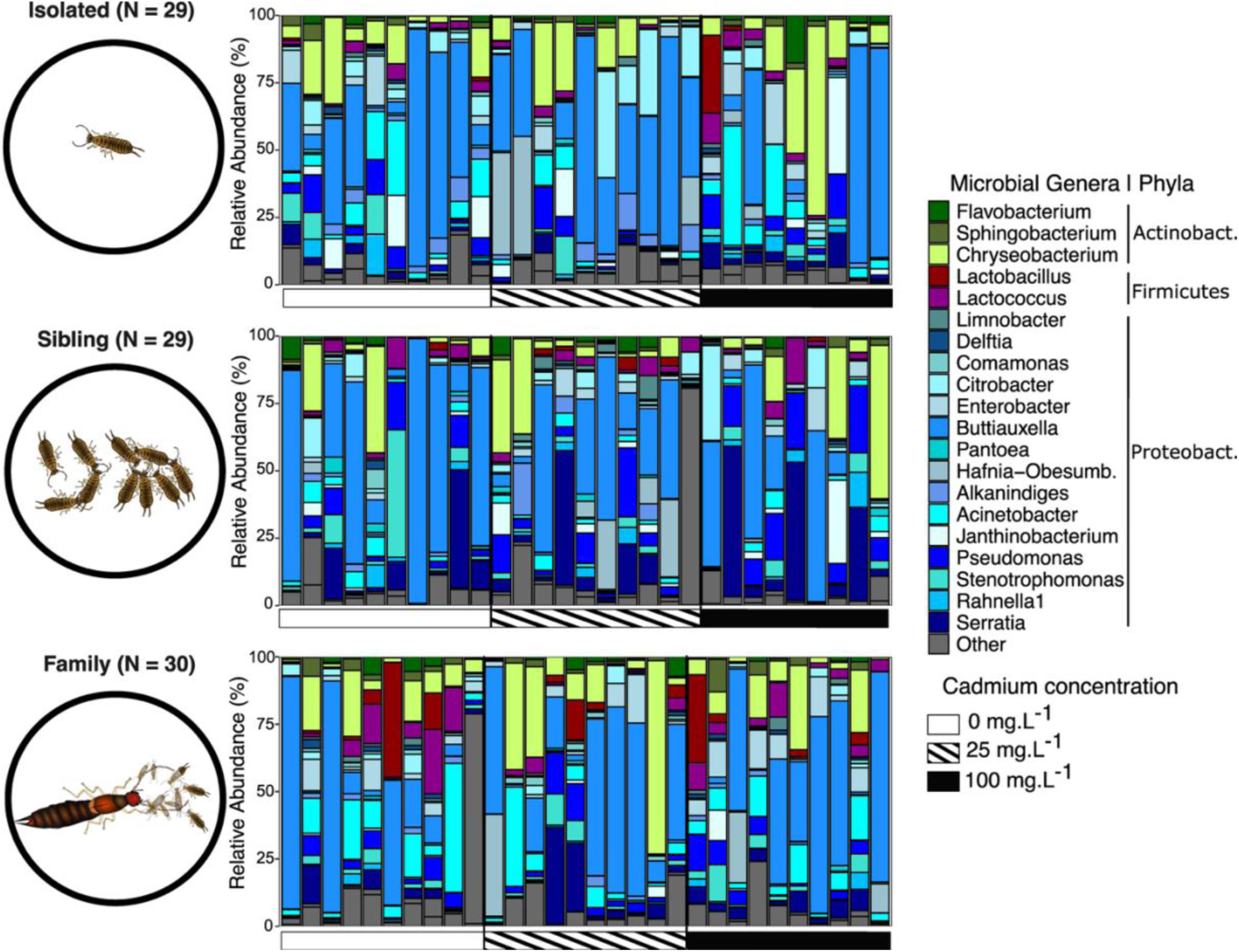
Microbiome composition of nymphs from isolated, sibling and family treatments. Barplots represent the individual composition for the top 20 microbial genera. Taxa belonging to Proteobacteria are in blue, Firmicutes in red/purple, and Bacteroidota in green. The gray contrasted horizontal bar represent the cadmium concentration for exposed nymphs. N indicates the number of nymphs used per social environment.

### Effects of social environment and cadmium on microbial diversity

Social environment and cadmium ingestion had independent effects on the unweighted Unifrac metric (Figure 2B), but no effect on alpha diversity (Figure 2A, Figure S4) neither the other beta diversity metrics (Figure S4). The social environment affected the unweighted Unifrac dissimilarities at both ASV level (R^2^ = 0.063, F = 2.90, P_adj_ = 0.016; Figure S4) and Genus level (R^2^ = 0.070, F = 3.28, P_adj_ = 0.016). This effect showed that nymphs exhibited microbiome structures that were specific of their social environment (Table 1, Figure 2B): isolated vs family treatments (ASV R^2^ = 0.066, F = 3.10, P_adj_ = 0.001; Genus R^2^ = 0.075, F = 4.61, P_adj_ = 0.001), isolated vs sibling treatments (ASV R^2^ = 0.039, F = 2.25, P_adj_ = 0.018; Genus R^2^ = 0.047, F = 2.73, P_adj_ = 0.022), and family vs sibling treatments (ASV R^2^ = 0.038, F = 2.25, P_adj_ = 0.014; Genus R^2^ = 0.038, F = 2.23, P_adj_ = 0.029). These results were consistent with the first two axes of the PCoA associated with the unweighted Unifrac distances where microbial structures of isolated nymphs were seemingly projected in the opposite direction of other samples along the first axis (Figure 2C). Microbial taxa such as bacteria belonging to the Phylum Bdellovibrionota and bacterial genera *Niabella* (Bacteroidota, Chitinophagaceae), *Rurimicrobium* (Bacteroidota, Chitinophagaceae), *Pedobacter* (Bacteroidota, Sphingobacteriaceae) or *Alkanindiges* (Proteobacteria, Moraxellaceae) were significantly correlated along the PCoA axes (P < 0.01) and tended to project in the direction of isolated nymph’s microbiomes along the first PCoA axis (Figure 2B).

**Figure 2.**
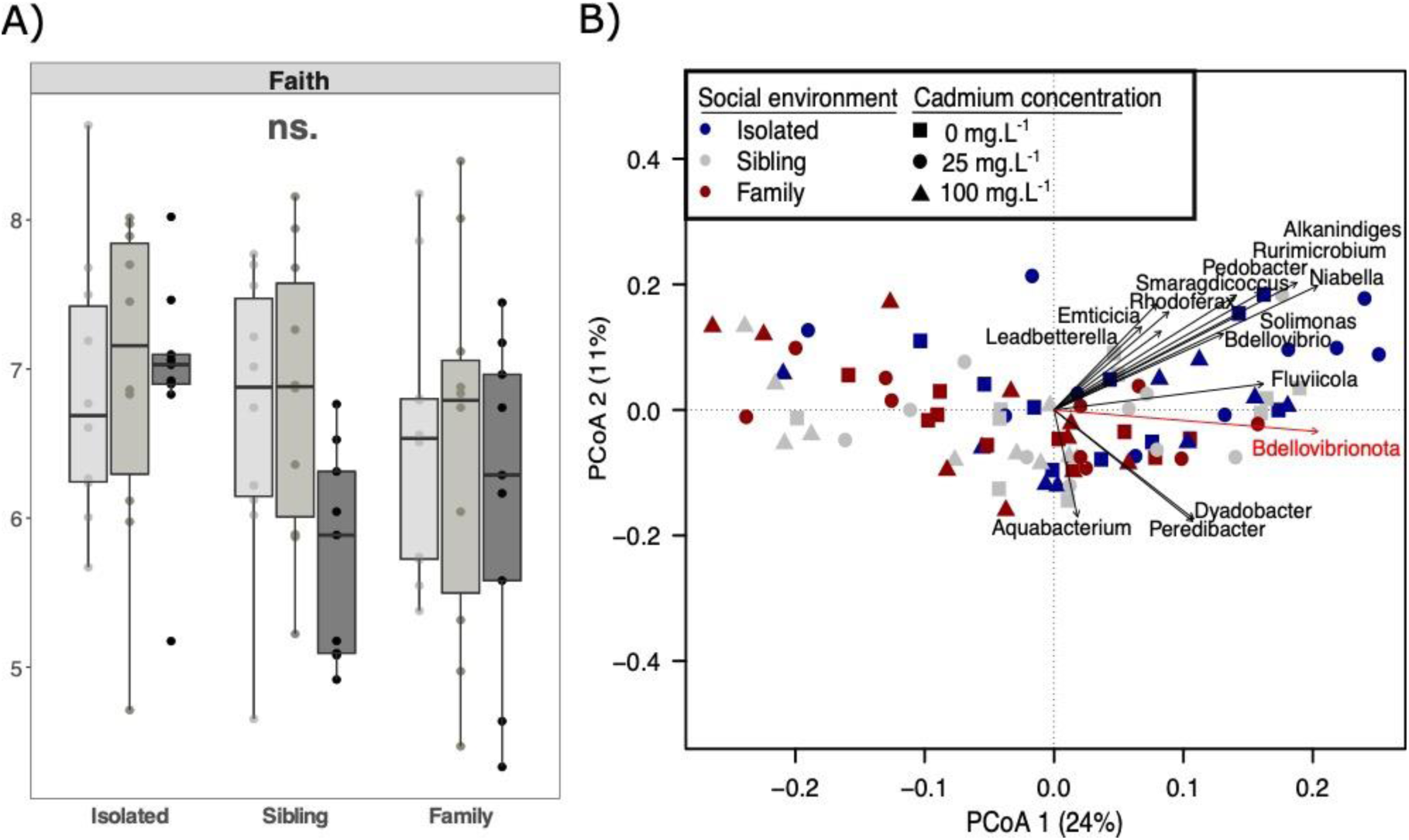
Phylogenetic alpha and beta-diversity of the nymph microbiomes. Differences in alpha-diversity where **(A)** the Faith (phylogenetic diversity) and **(B)** the unweighted Unifrac distances of the ASVs composition between samples were tested according to the social environment and the cadmium concentration. Boxplots show the median (middle bar) and the interquartile range (box) with whiskers representing the 1.5-fold the interquartile range. Non-significant results (Padj > 0.05) are labelled by ‘ns’. Dissimilarities are visualized by the first two axes of a Principal Coordinates Analyses (PCoA), where each nymph is coloured according to their social environment and shaped by the cadmium concentration. The microbial genera and phyla that are significantly correlated (P < 0.01) are projected in black and red, respectively. Additionally, cadmium ingestion marginally affected unweighted Unifrac dissimilarities at ASV level (R^2^ = 0.041, F = 1.88, P_adj_ = 0.053). Nymphs exposed to 100 mg.L^-1^ cadmium had a different microbial structure than nymphs exposed to 0 mg.L^-1^ (R^2^ = 0.035, F = 2.04, P_adj_ = 0.023) and 25 mg.L^-1^ (R^2^ = 0.038, F = 2.21, P_adj_ = 0.018), whereas there was no difference between 0 mg.L^-1^ and 25 mg.L^-1^ (R^2^ = 0.022, F = 1.29, P_adj_ = 0.188). Note, however, that these last results must be taken with caution as dispersion (*i.e.,* homogeneity) of microbiomes regarding the unweighted Unifrac distances differed between the three cadmium concentrations with highest distance from the centroid in the concentration 25 mg.L^-1^ (P_adj_ = 0.026). Finally, we did not detect any effect of the interaction between social environment and cadmium ingestion, either for alpha diversity (all P_adj_ ≥ 0.136) or beta diversity (all P_adj_ ≥ 0.169; Table 1).

**Table 1.**
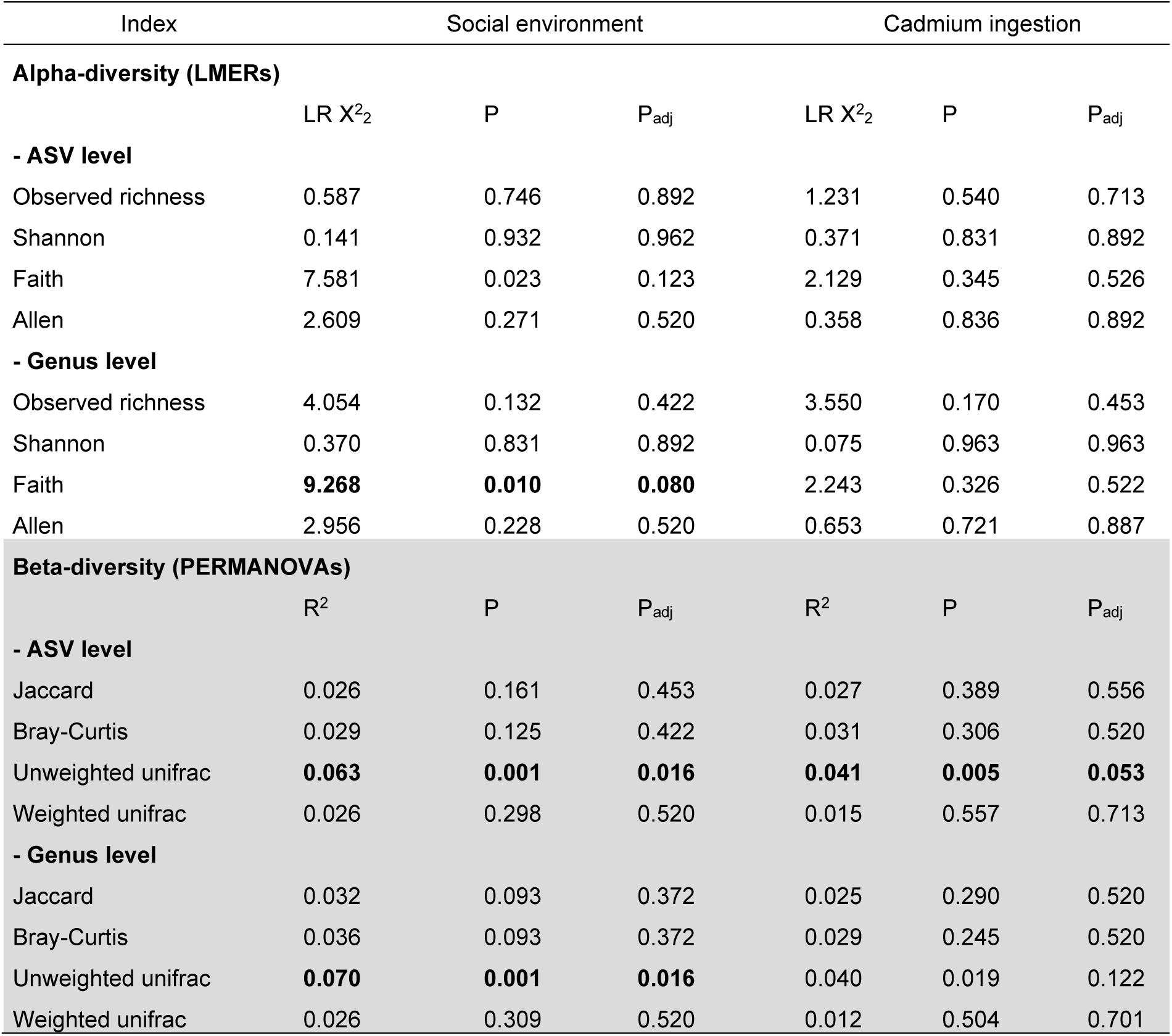
Effects of social environment, cadmium ingestion, and their interaction on the alpha-diversity and beta-diversity of nymph microbiomes at both ASV and Genus levels. Statistical values from linear models with random effects (LMERs) or PERMANOVAs conducted on each index. P-values were corrected for multiple testing according to False Discovery Rate (FDR) correction. The models were simplified by removing the interaction as it was not significant for both alpha (all Padj ≥ 0.136) and beta diversity (all Padj ≥ 0.169). Significant results (Padj ≤ 0.05) are indicated in bold.

### Effect of social environment and cadmium on bacterial genera

Out of a total of 84 bacterial genera that formed the nymph’s microbiome, 18 and 7 were statistical indicators of the social environment and the cadmium concentration, respectively. Of these, 13 and 5 bacterial genera showed a |log2fold change| > 2 (*i.e.,* for which the abundance is at least 4 times different), respectively. Among these bacteria, *Janthinobacterium* (Proteobacteria, Oxalopacteraceae), *Kosakonia* (Proteobacteria, Enterobacteriaceae), *Pedobacter* (Bacteroidota, Sphingobacteriaceae), *Rurimicrobium* (Bacteroidota, Chitinophagaceae) were more abundant in isolated nymphs than in sibling or family nymphs (Figure 3; Table S1). In contrast, the bacterial genera *Duganella* (Proteobacteria, Oxalobacteraceae), *Cellvibrio* (Proteobacteria, Cellvibrionaceae) and *Serratia* (Proteobacteria, Yersiniaceae) dominated the sibling nymphs. *Rhodococcus* (Actinobacteria, Nocardiaceae), *Lactobacillus* (Firmicutes, Lactobacillaceae), *Paracoccus* (Proteobacteria, Rhodobacteraceae), *Tsukamurella* (Actinobacteria, Tsukamurellaceae), *Erwinia* (Proteobateria, Erwiniaceae) and *Sphingobium* (Proteobacteria, Sphingomonadaceae) were the most abundant bacteria in family nymphs (Figure 3; Table S1).

**Figure 3.**
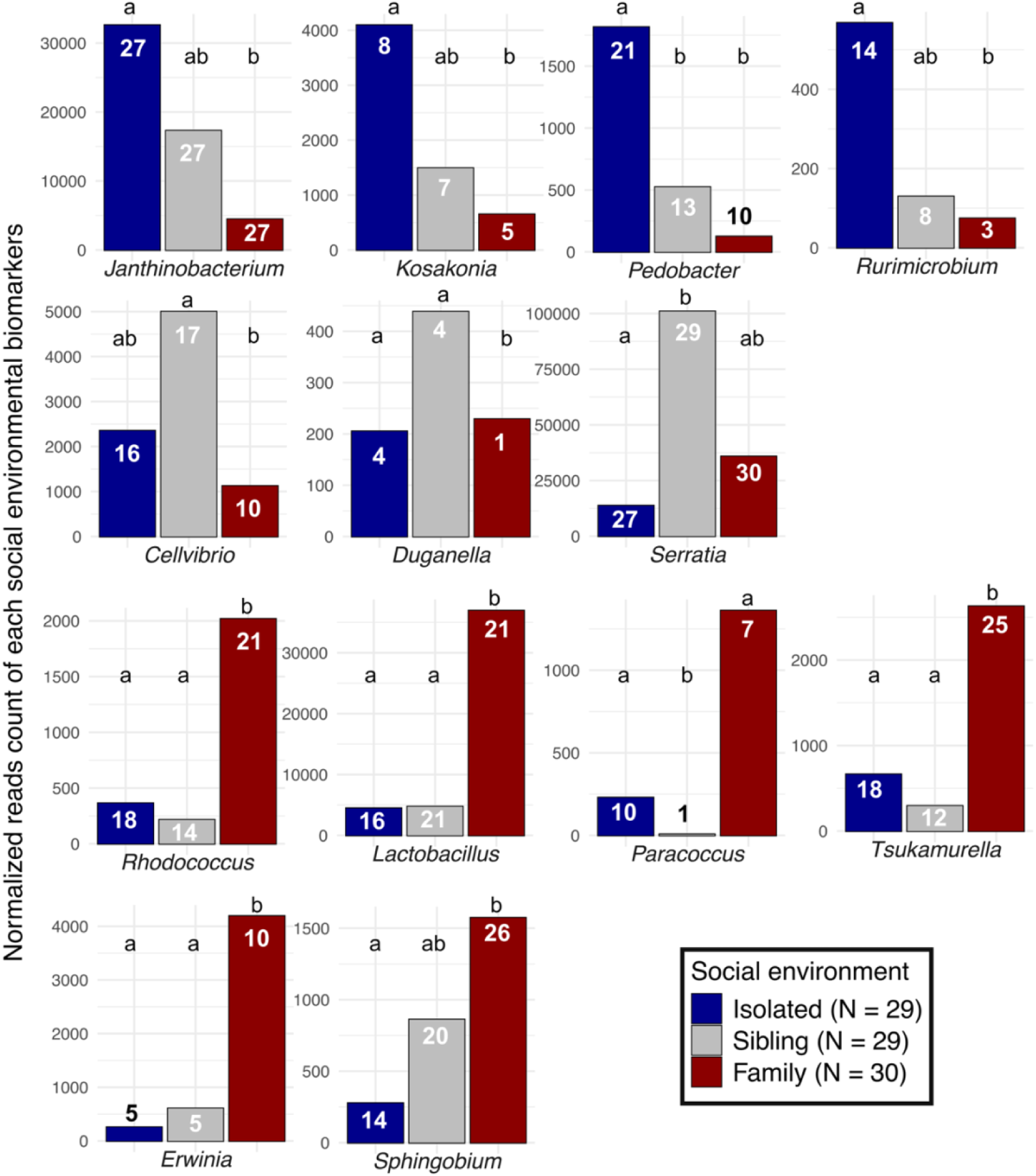
Normalized reads count of each bacterial genus indicators related to isolated, sibling or family nymphs. Indicators are delineated using a DESeq2 approach (P < 0.05) for which the significant contrast showed a |log2fold change| > 2. The prevalence for each genus is labelled in the bar of the corresponding social environment. Different letters indicate P < 0.05.

Regarding cadmium ingestion, we detected a striking increase in *Duganella* (Proteobacteria, Oxalobacteraceae), *Hafnia-Obesumbacterium* (Proteobacteria, Hafniaceae), *Erwinia* (Proteobateria, Erwiniaceae), *Nubsella*, *Pedobacter* (Bacteroidota, Sphingobacteriaceae) and *Rurimicrobium* (Bacteroidota, Chitinophagaceae) in nymphs exposed to 25 mg.L^-1^ of Cd compared to nymphs exposed to 0 or 100 mg.L^-1^ of Cd (Figure 4; Table S1). None of these bacteria were represented as discriminant of both parameters (*i.e.,* social environment and cadmium ingestion), suggesting that the effect of the social environment does not influence (or is not influenced) by the cadmium ingestion.

**Figure 4.**
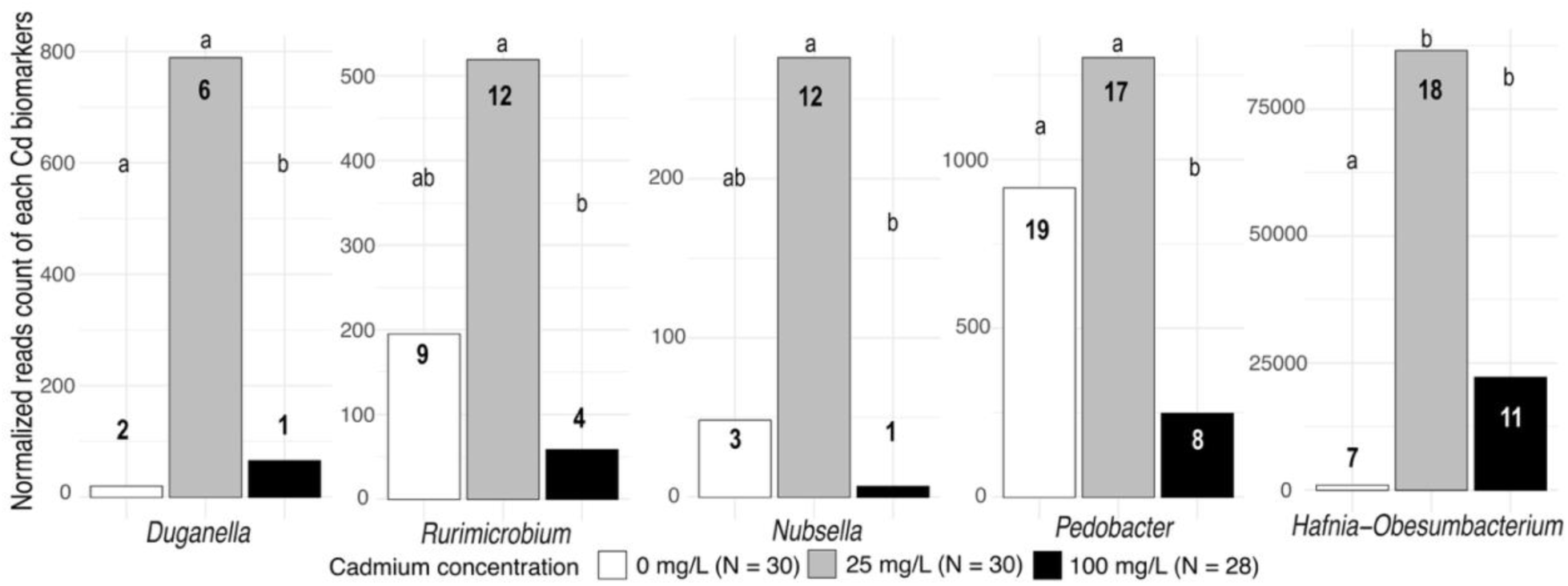
Normalized reads count of each bacterial genus indicators related to cadmium concentration. Indicators are delineated using a DESeq2 approach (P < 0.05) for which the significant contrast showed a |log2fold change| > 2. The prevalence for each genus is labelled in the bar of the corresponding cadmium concentration. Different letters indicate P < 0.05.

## Discussion

Understanding the factors that drive changes in communities of mutualistic and commensal microorganisms in animal hosts is crucial for enhancing our knowledge of host fitness, population dynamics and ecosystem functioning (Martiny et al., 2023). This is particularly true in the context of anthropogenic environmental changes, which are increasingly known to both directly and indirectly shape host-symbiont relationships (Cheutin et al., 2021; Maraci et al., 2022; Teyssier et al., 2018). In this study, we tested the simultaneous effects of the social environment and exposure to a chemical pollutant on the diversity and nature of the microbial communities present in earwig juveniles. Our results first show that the social environment affected the phylogenetic diversity of the microbial communities (*i.e.,* unweighted Unifrac distances) at both the bacterial sequence (*i.e.,* ASV) and genus levels. Our results then show that cadmium ingestion influenced the phylogenetic qualitative composition of the microbiome at the ASV level. Notably, we found that changes in the microbiome induced by the social environment and cadmium ingestion were associated with the turnover of specific bacterial taxa. However, there was no effect of the social environment and cadmium ingestion on the other indices of alpha-diversity and beta-diversity of the nymph microbiome. Finally, as we found no interaction between cadmium and social environment, our results suggest that cadmium ingestion did not affect the social transmission of the microbiome to earwig nymphs.

Our first finding is that nymphs exhibited social environment-dependent microbial communities according to the unweighted Unifrac metric. This indicates that nymphs reared alone, with siblings, or with a full family group (siblings plus mother) harbour distinct microbiome profiles, each primarily shaped by the rare or low abundance of bacteria from phylogenetically distant taxa. These results provide two key insights. First, the presence of specific taxa in the isolated nymphs suggest that living in a group may not only enhance access to new bacterial sources, as reported in other social mammals and insects (Hosokawa et al., 2007; Raulo et al., 2021; Tung et al., 2015; Y. Wang & Rozen, 2017), but also help to prevent colonization by opportunistic environmental taxa. This is examplified by the phylum Bdellovibrionota (including the two genera *Peredibacter* and *Bdellovibiro*), which was more abundant in isolated nymphs compared to those reared with siblings or within a full family group (Figure 2B). This phylum is typically abundant in the soil and is occasionally found in the gut microbiome of certain insects (Davidov et al., 2006). The proliferation of Bdellovibrionota in isolated nymphs might thus stem from their initial acquisition from the environment, combined with the absence of competitive bacterial strains that are typically transmitted by family members (Figueiredo & Kramer, 2020). Further testing is needed to validate this hypothesis. The second key finding is that the composition of the group plays an important role in shaping the nymph’s microbiome. Some bacterial taxa were more abundant in nymphs reared in a whole-family group than in nymphs reared with their siblings only. This suggests a maternal transmission of bacteria to nymphs during earwig family life. Interestingly, many of the bacteria putatively transmitted by earwig mothers have known beneficial functions related to digestion and immunity in other (social) insects. For example, the Proteobacteria *Sphingobium* or *Rhodococcus* present a large variety of catabolic functions such as aromatic hydrocarbons metabolization or cellulose digestion in cockroaches, termites or beetles (Biswas & Paul, 2021; Delalibera et al., 2005; Wenzel et al., 2002). Similarly, the bacterium *Lactobacillus* can protect against the negative effects of pathogens, pesticides and cadmium, as well as help for the fermentation of carbohydrate plants and cereals in bees, honeybees or grasshoppers (Lazzeri et al., 2020; Li et al., 2021; Nowak et al., 2021; Vásquez et al., 2012). To date, the role of the gut microbiome in the fitness of the European earwig remains unclear (Cheutin, Leclerc, et al., 2024; Van Meyel et al., 2021), and future experimental and transcriptomic studies are needed to investigate the potential function of these bacteria in this species.

We found that the social environment affected the nymph microbiome not only through differences in rare/low-abundant phylogenetically distant taxa, but also via an over-representation of 13 bacterial taxa (4 specific to the isolated treatment, 3 to the sibling treatment, and 7 to the full family treatment). In isolated nymphs, two of these four taxa (*Kosakonia* and *Janthinobacterium)* are insect mutualists and typically enhance host protection against pathogens. For instance, *Janthinobacterium* has antifungal functions in many insects (Haack et al., 2016; Magoga et al., 2023; Rodrigues et al., 2006), while *Kosakonia* can inhibit the growth of the entomopathogenic strains belonging to the bacteria *Serratia* (Grimont et al., 1979; Muratoğlu et al., 2011) and render mosquitoes and tsetse flies hosts refractory to parasite infections such as *Trypanosoma* and *Plasmodium* (Weiss et al., 2019). The overabundance of these bacteria in isolated nymphs might be an adaptive strategy, enabling them to mount an effective defense against pathogens in the absence of social immunity provided by siblings and mothers (Diehl et al., 2015; Diehl & Meunier, 2018). This hypothesis would also be consistent with our results showing that *Serratia* (an opportunistic entomopathogen) is less abundant in isolated than group-living nymphs (see above). Independent of the social environment, we found that cadmium exposure also played an important role in shaping the diversity of the nymph microbiome. First, cadmium ingestion induced unstable community states consistent with the Anna Karenina principle (Zaneveld et al., 2017), whereby microbiomes became more heterogeneous and compositionally distinct in nymphs exposed to the highest (100mg.L^-1^) compared to intermediate (25 mg.L^-1^) or control (0 mg.L^-1^) doses, as revealed by both the Faith and unweighted Unifrac metrics. This effect is in line with the typical effects of cadmium on microbial communities reported in a taxonomically wide range of animals (Li et al., 2021; Rothman et al., 2019; Trevors et al., 1986; S. Zhang et al., 2015; Y. Zhang et al., 2016). Second, cadmium ingestion at the intermediate concentration was associated with the presence or absence of five bacterial taxa. Of these, only one (*Hafnia-Obesumbacterium*) has been reported in animal microbiomes (S. Zhang et al., 2023). This bacterium is a pathogen in many species, where its abundance often increases under stressful conditions (Janda & Abbott, 2006). For example, *Hafnia-Obesumbacterium* is dominant in the gut microbiomes of tadpole *Rana chensinensis* and crayfish *Procambarus clarkii*, respectively exposed to lead pollution (S. Zhang et al., 2023) and acidic conditions (Z. Wang et al., 2024). As such, our data suggest that this genus is the only bacterium sensitive to cadmium ingestion in juveniles of the European earwig. The other four bacterial taxa specific to the intermediate dose of cadmium probably reflect the good development of these bacteria on medium contaminated with low levels of cadmium and their subsequent passive environmental uptake by earwig nymphs via contact with cadmium-contaminated food. This is consistent with previous studies indicating that the growth of *Pedobacter* and *Duganella* is typically favoured in cadmium-enriched soils (X. Cheng et al., 2022; Stout et al., 2010; H. Wang et al., 2023). However, while our findings align with the existing literature, they must be considered with caution, as they are based on metabarcoding approach (Djemiel et al., 2022). Given the qualitative nature of our results, they may be biased due to their compositional framework. Quantitative and transcriptomic studies are now needed to validate these findings and further elucidate the functional role of the microbiome under environmental stress (Gloor et al., 2017).

Overall, this study shows that the access to a social environment, the composition of family group and exposure to chemical pollutants can all influence the composition and diversity of the microbiome in juvenile European earwigs. Interestingly, we show that these effects do not interact with each other, *i.e.* varying the access to a social environment or changing the composition of that nymph microbiome, and vice versa. As anthropogenic environmental changes continue to accelerate globally (Johnson & Munshi-South, 2017; Seto et al., 2012), these results demonstrate that the ability to naturally switch between solitary and social life (as found in the European earwig nymphs, Meunier, 2024) does not necessarily confer an advantage in mitigating the negative effects of chemical contaminants (at least) on the host microbiome. Future studies are now needed to better understand the role of the earwig microbiome on its fitness and population dynamics (Cheutin, Leclerc, et al., 2024; Van Meyel et al., 2021), and to determine whether the reported effects of social environment and cadmium intake on its diversity can alter these parameters.

## Data availability

The data set and the scripts are available on Zenodo with the url https://zenodo.org/doi/10.5281/zenodo.13380552. The reads for each sample were deposited in the NCBI Sequence Read Archive (SRA) under the BioProject accession no. PRJNA1152783.

## Acknowledgements

We thank Charlotte Lécureuil for her comments on this manuscript. We also thank all members of the EARWIG group at IRBI for their help with animal rearing and setup installation, and the Bio-environment platform of the University of Perpignan (France) for providing the MiSeq Illumina. We thank Armand Guillermin and Stéphanie Drusch of the INRAE unité expérimentale Recherche Intégrée in Gotheron for giving us access to their orchards for earwig field sampling. The graphical abstract was made with BioRender application. Preprint version 3 of this article has been peer-reviewed and recommended by Peer Community In Microbiology ( https://doi.org/10.24072/pci.microbiol.100161; Cheutin, Honorio and Meunier, 2025. Finally, we also thank the two anonymous reviewers for their insightful comments in a previous version of this manuscript and the precious recommendation of Thomas Pollet.

## Author’s contribution

**Marie-Charlotte Cheutin**: Data curation; Formal analysis; Investigation; Methodology; Writing - original draft; and Writing - review & editing. – **Romain Honorio**: Conceptualization; Methodology; Writing - original draft; and Writing - review & editing. **Joël Meunier**: Conceptualization; Funding acquisition; Supervision; Writing - review & editing

## Funding information

This work was funded by the Région Centre-Val de Loire (Project DisruptCare; n°2021 00149474) and by a research grant from the Agence Nationale de la Recherche (ANR-20-CE02-0002).

## Conflict of interest

The authors declare that this study had no conflicts of interest.

## Ethics approval statements

Our investigation complies with the current European Directive 2010/63/EU that does not acquire ethical approval on invertebrates. All animals were handled with care until necessary sacrifices.

## Notes

https://zenodo.org/doi/10.5281/zenodo.13380552

### Competing Interest Statement

The authors have declared no competing interest.

### Summary of Updates

This new version has been recommended by the peer community PCI Microbiology. We added add a hyperlink on the badge pointing to https://doi.org/10.24072/pci.microbiol.100161.

