## Supplementary material for "Family life and cadmium ingestion independently shape offspring microbiomes in a subsocial insect": Table S1, Figure S1, Figure S2, Figure S3, Figure S4

Running title : No cadmium effect on juvenile earwigs' microbiome

Marie-Charlotte Cheutin*^1^, Romain Honorio*^1,2^ & Joël Meunier^1^

^1^ Institut de Recherche sur la Biologie de l’Insecte, UMR 7261, CNRS, University of Tours, Tours, France

^2^ Laboratoire d’Ethologie Expérimentale et Comparée UR 4443 (LEEC), Université Sorbonne Paris Nord, 93430 Villetaneuse, France

* These authors contributed equally to this work

ORCID

MC Cheutin #0000-0001-7711-7512

R Honorio #0000-0001-7623-8427

J Meunier #0000-0001-6893-2064

**Table S1: Results of the DESeq2 analysis regarding the effects of the cadmium ingestion and the social environment.**

| **Genus** | **Family** | **Phylum** | **Log2FC** | **P** | **Contrast** |
| --- | --- | --- | --- | --- | --- |
| **Social environment contrasts** | | | | | |
| Rhodococcus | Nocardiaceae | Actinobacteriota | 2,29 | .019 | Family - Isolated |
|  |  |  | 2,23 | .028 | Family - Sibling |
| Tsukamurella | Tsukamurellaceae |  | 2,01 | .041 | Family - Isolated |
|  |  |  | 2,90 | .002 | Family - Sibling |
| Pedobacter | Sphingobacteriaceae | Bacteroidota | -4,51 | > .001 | Family - Isolated |
|  |  |  | -2,89 | .013 | Sibling - Isolated |
| Rurimicrobium | Chitinophagaceae |  | -4,01 | .014 | Family - Isolated |
| Lactobacillus | Lactobacillaceae | Firmicutes | 3,54 | .001 | Family - Isolated |
| Lactococcus | Streptococcaceae |  | 1,39 | .021 | Sibling - Isolated |
| Acinetobacter | Moraxellaceae | Proteobacteria | 1,41 | .001 | Family - Sibling |
| Alkanindiges |  |  | -1,44 | .001 | Family - Isolated |
|  |  |  | -1,34 | .007 | Sibling - Isolated |
| Cellvibrio | Cellvibrionaceae |  | -3,11 | .047 | Family - Sibling |
| Duganella | Oxalobacteraceae |  | -23,46 | > .001 | Family - Isolated |
|  |  |  | -2.86 | > .001 | Family - Sibling |
| Erwinia | Erwiniaceae |  | 8,83 | > .001 | Family - Isolated |
|  |  |  | 5,51 | .028 | Family - Sibling |
| Janthinobacterium | Oxalobacteraceae |  | -3,09 | > .001 | Family - Isolated |
|  |  |  | -1,80 | .035 | Family - Sibling |
| Kosakonia | Enterobacteriaceae |  | -5,77 | .028 | Family - Isolated |
| Paracoccus | Rhodobacteraceae |  | 7,86 | .001 | Family - Sibling |
|  |  |  | -6,08 | .021 | Sibling - Isolated |
| Pseudomonas | Pseudomonadaceae |  | -1,21 | .035 | Family - Sibling |
|  |  |  | 1,40 | .021 | Sibling - Isolated |
| Rahnella1 | Yersiniaceae |  | -1,95 | .016 | Family - Sibling |
| Serratia |  |  | 1,53 | .027 | Family - Isolated |
|  |  |  | 2,75 | > .001 | Sibling - Isolated |
| Sphingobium | Sphingomonadaceae |  | 2,47 | .002 | Family - Isolated |
| **Cadmium exposition contrasts** | | | | | |
| Nubsella | Sphingobacteriaceae | Bacteroidota | -5,22 | .022 | 100 - 25 |
| Pedobacter |  |  | -3,32 | .001 | 100 - 0 |
|  |  |  | -3,30 | .002 | 100 - 25 |
| Rurimicrobium | Chitinophagaceae |  | -4,12 | .022 | 100 - 25 |
| Alkanindiges | Moraxellaceae | Proteobacteria | -1,49 | .001 | 100 - 0 |
|  |  |  | -1,82 | > .001 | 100 - 25 |
| Duganella | Oxalobacteraceae |  | -25,05 | > .001 | 100 - 0 |
|  |  |  | -32,00 | > .001 | 100 - 25 |
| Enterobacter | Enterobacteriaceae |  | 1,39 | .038 | 100 - 0 |
| Erwinia | Erwiniaceae |  | -5,82 | .030 | 100 - 0 |
| Hafnia-Obesumbacterium | Hafniaceae |  | 6,44 | > .001 | 100 - 0 |
|  |  |  | 9,60 | > .001 | 25 - 0 |
| Rahnella1 | Yersiniaceae |  | 1,88 | .022 | 100 - 25 |

**Figure S1: Electrophoresis gel (agarose 1.5%) of the 16S rRNA amplicons.** Samples were amplified using the specific prokaryotic probes 343F and 784R coupled with Illumina linkers (amplicon product ~ 500 bp). Two extraction blanks (*i.e.,* dissection tools washed in T1 buffer and one empty tube filled with T1 buffer) at the begin of the PCR plate, and one PCR blank filled with molecular water at the end of the plate, were also amplified and did not amplify (marked as -). One extraction and one PCR positive amplicon (marked as +) were realized with one sample that had been already successfully sequenced. The success of the remaining samples was revealed in another electrophoresis gel.

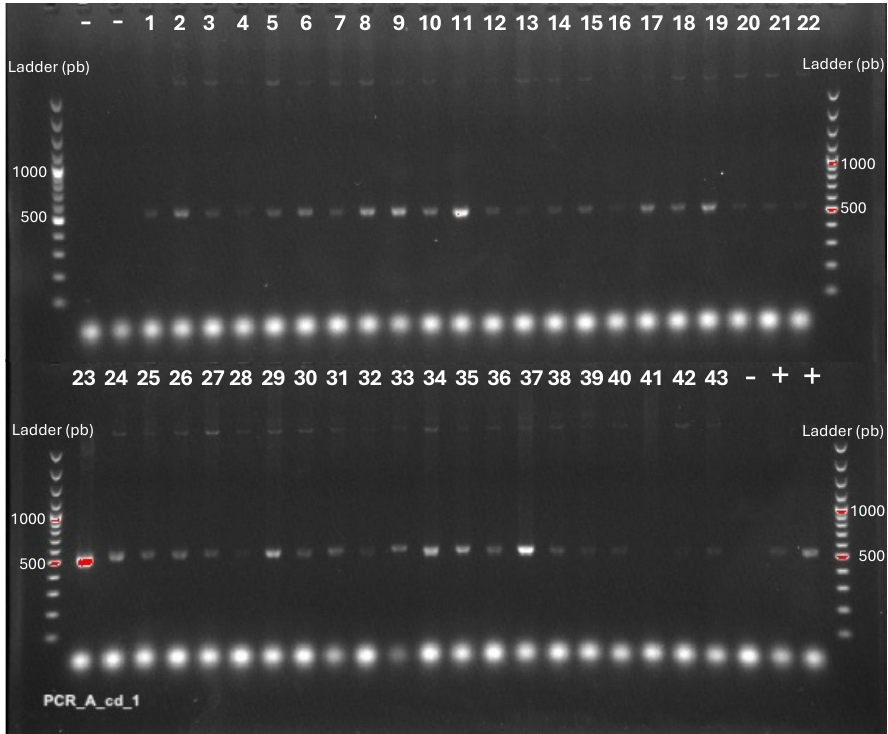

**Figure S2: Species-Abundance Distribution (SAD) patterns for the microbial ASVs of the European earwig nymphs.** Each ASV (coloured according to its Phylum) is plotted according to its dispersion index and its occurrence in all samples. The curve indicates the threshold above which ASVs are under-dispersed compared to Poisson distribution (*i.e.,* not random) and are considered as core ASVs, while those below are considered as clustered and correspond to satellite ASVs.

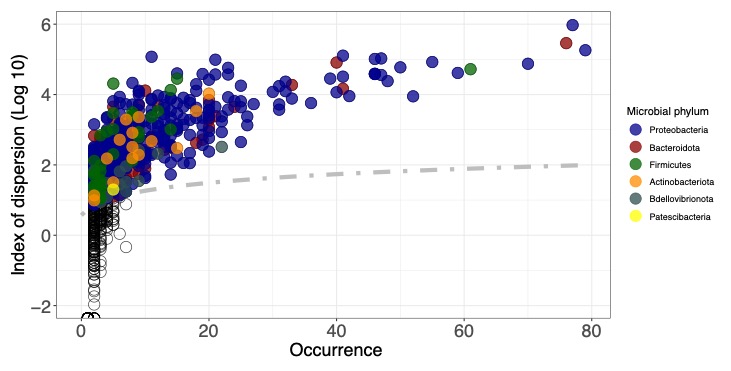

**Figure S3: Rarefaction curves for each nymph microbiome, depending on its social environment.**

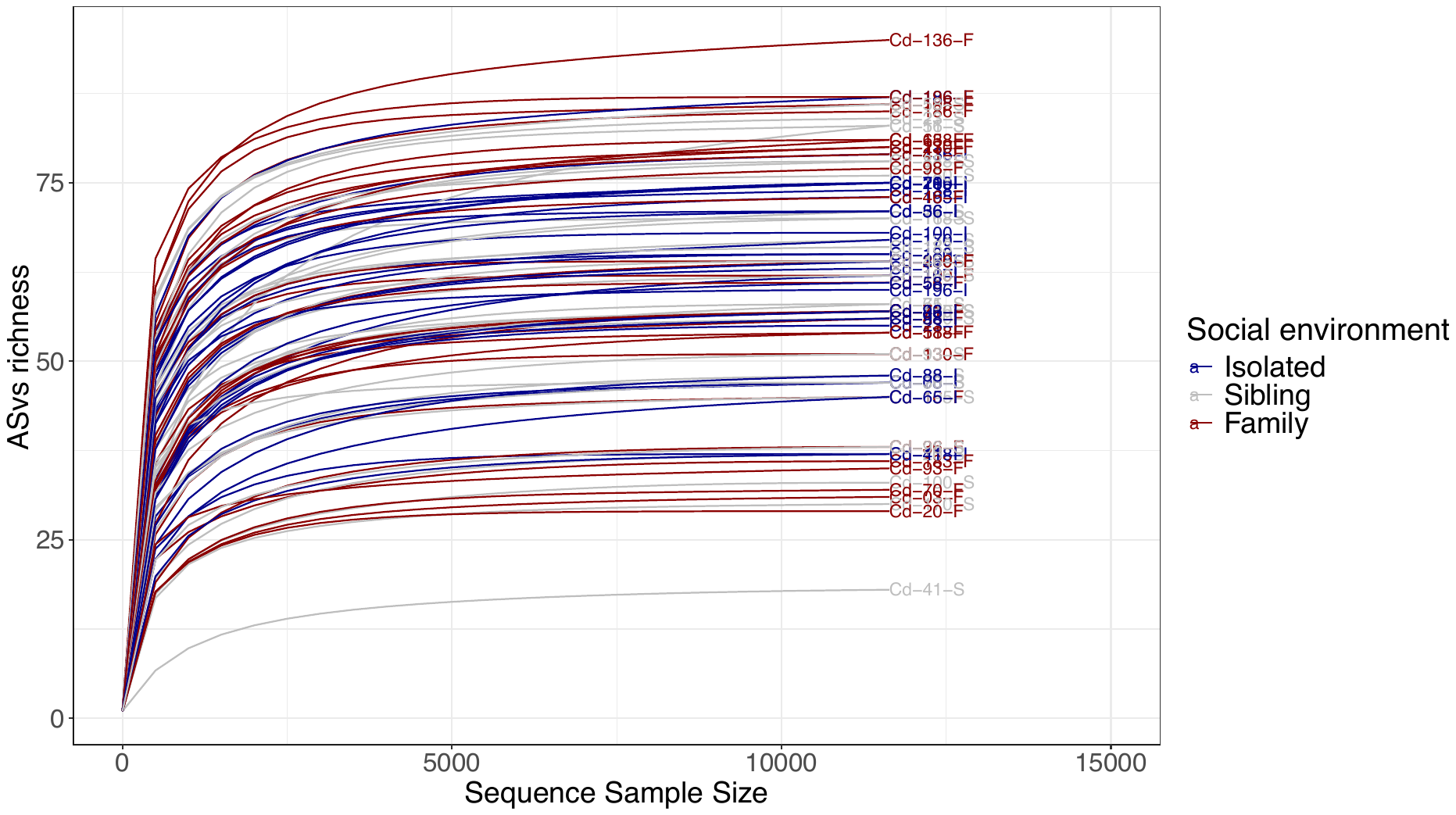

**Figure S4: Alpha and beta-diversity** **of the nymph microbiomes.** Differences in alpha-diversity where **(A)** the Observed richness, **(B)** the Shannon and the **(C)** Allen indices were tested according to the social environment and the cadmium concentration. Boxplots show the median (middle bar) and the interquartile range (box) with whiskers representing the 1.5-fold the interquartile range. Non-significant results (P_adj_ > 0.05) are labelled by ‘ns’. Beta-diversity is based on the (**D**) Jaccard, (**E**) Bray-Curtis and (**F**) weighted Unifrac distances of the ASVs composition between samples. Dissimilarities are visualized by the first two axes of a Principal Coordinates Analyses (PCoA), where each nymph is coloured according to their social environment and shaped by the cadmium concentration. The microbial genera and phyla that are significantly correlated (P < 0.01) are projected in black and red, respectively.
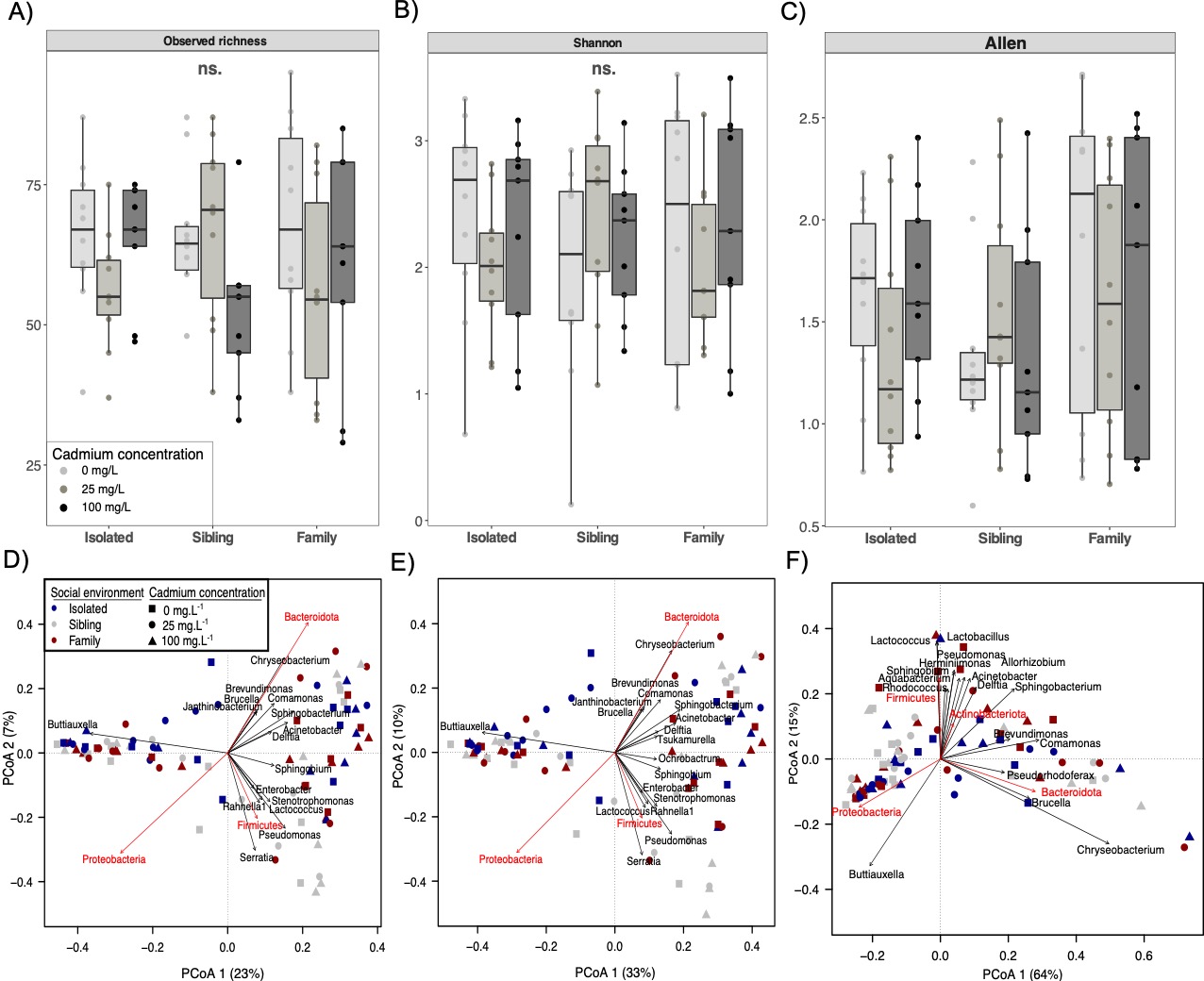
